# An amphipathic and cationic antimicrobial peptide kills colistin resistant Gram-negative pathogens *in vivo*

**DOI:** 10.1101/2021.12.13.472350

**Authors:** Thomas T. Thomsen, Mette Kolpen, Vinoth Wigneswaran, Ulrik Kromann, Anna E. Ebbensgaard, Anette M. Hammerum, Henrik Hasman, Stine Radmer, Kasper N. Kragh, Rasmus Hartmann-Petersen, Paul Robert Hansen, Anders Folkesson, Niels Frimodt-Møller, Anders Løbner-Olesen

## Abstract

New antibiotics are needed against multidrug resistant Gram-negative pathogens that have compromised global health systems. Antimicrobial peptides are generally considered promising lead candidates for the next generation of antibiotics but have not fulfilled this expectation. Here we demonstrate activity of a cationic amphipathic undecapeptide (ChIP; <u>Ch</u>arge change <u>I</u>ndependent <u>P</u>eptide) against a wide panel of multidrug resistant Gram-negative pathogens. Importantly, the antimicrobial activity of ChIP is independent of the surface charge changes that confer colistin resistance through modification of Lipid A, while decreased activity of ChIP correlates with GlcN1 tri-acylation of Lipid A. In an *in vivo* peritonitis mouse model ChIP displays excellent activity against both colistin sensitive and resistant *Escherichia coli* and *Acinetobacter baumannii* strains.

**Author Summary:** Antimicrobial peptides hold promise as novel treatment options for diseases caused by multidrug resistant bacteria. Here we present evidence that the ChIP peptide, comprised of 11 D-amino acids, is active against a variety of Gram-negative bacteria that ranks high on the WHO list of critically important pathogens. ChIP initially interacts with the Gram-negative outer membrane, independent of its surface charge, followed by entry into the periplasm and permeabilization of the inner membrane, leading to bacterial cell death. Detailed analyses of the outer membrane indicate that the acylation pattern of lipopolysaccharides plays an important role for ChIP activity. In a mouse infection model, ChIP display excellent activity in reducing bacterial numbers for both *Escherichia coli* and *Acinetobacter baumannii*. Importantly, ChIP is highly efficient against bacteria resistant to colistin, an antibiotic normally considered as a last resort treatment of infections caused by multidrug resistant Gram-negative bacteria.

## Introduction

During the last decades, infections caused by multidrug resistance (MDR) bacteria has become a major global health concern, as antimicrobial therapy options dwindle, and the development of new antimicrobial agents has stagnated (1). Special concerns have been raised for MDR species from the ESKAPE group (*Enterococcus faecium, Staphylococcus aureus, Klebsiella pneumoniae, Acinetobacter baumannii, Pseudomonas aeruginosa and Enterobacter species*) (1, 2). Therefore, it is necessary to search for and develop novel treatment options for these ESKAPE pathogens. Among the possibilities for antimicrobial development, antimicrobial peptides (AMP) have gained extensive interest (3-5). AMP exist as part of the natural immune defense of most multicellular organisms (4, 5). Generally, AMP consist of 20-40 amino acid residues, act on the bacterial membrane and are active at low concentrations against a wide range of microorganisms (4, 5). While AMP are highly modifiable, there is a lack in understanding of their mode of action and specificity to bacterial membranes. The main caveats of AMP based antibiotics often relates to toxicity, expensive manufacturing cost and low bioavailability (6). However, such shortcomings may be overcome by AMP modification, including incorporation of non-natural amino acids, such as D-amino acids or peptidomimetics derivatives (7). AMP remain attractive for antimicrobial development, especially those targeting the bacterial envelope and not a single intracellular target, thereby circumventing the need for cellular uptake through the highly selective bacterial envelope. This mechanism of action is assumed to restrict AMP resistance development due to the genetic complexity in remodeling of the bacterial envelope (8, 9).

The envelope of Gram-negative bacteria is a formidable barrier evolved to protect the bacterial cell from a variety of extracellular stresses. The envelope consists of three distinct layers conferring protection. The innermost plasma membrane is composed of a phospholipid bilayer enclosing the intracellular environment. Encompassing the plasma membrane, the periplasmic space holds the cell stabilizing peptidoglycan which is attached to the first line defense, the outer membrane (OM) (10). The OM is an asymmetric bilayer with an inner leaflet of phospholipids and an outer leaflet composed almost entirely of lipopolysaccharide (LPS) (11, 12). LPS consist of three distinct regions: the membrane-anchoring Lipid A, the inner core and the outer core to which the O-antigen is attached (10). The negatively charged LPS which protrudes from the cell, is stabilized laterally by cations (Ca^2+^ and Mg^2+^), creating a highly impenetrable barrier to protect the cell from toxic compounds such as detergents and hydrophobic antibiotics (10).

The cationic lipopeptide colistin (polymyxin E) (13), is generally considered a last resort antibiotic against infections caused by MDR Gram-negative pathogens. Structurally, colistin consists of a positively charged cyclic peptide to which a lipophilic tail is coupled via an exocyclic peptide. Colistin, like many other AMP, targets the Gram-negative membrane. Initially colistin interacts with LPS in a charge-dependent manner, i.e. by electrostatic interactions, leading to destabilization of the cationic (Mg^2+^ and Ca^2+^) interactions between LPS molecules, before complete destabilization of the outer membrane by insertion of the lipophilic tail into the membrane (14). Until recently, the killing of the cells was described through this destabilization of the outer membrane, self-promoted uptake, and finally lysis of the inner membrane. A recent study has demonstrated that it is in fact the targeting of LPS at the plasma membrane that leads to lysis of the plasma membrane (15). For decades, colistin resistance development seemed unlikely due to its complex multimodal antimicrobial activity (3, 9). However, plasmid carried colistin resistance genes have recently disseminated across the globe (16, 17). These mobile colistin resistance genes named *mcr-1*–*9* (18), encodes phosphoethanolamine transferases that modify LPS with phosphoethanolamine (pEtN) moieties on Lipid A phosphates. This modification is said to lower the negative charge of the membrane, thereby diminishing affinity for colistin (11). Colistin resistance in Gram-negative species can also occur via constitutive activation of the genome encoded two-component pathways PhoP-PhoQ and BasS-BasR (also known as PmrA-PmrB), that regulate the downstream activation of LPS modifying enzymes EptA and ArnT (19-22). EptA is a phosphoethanolamine transferase similar to the enzymes encoded by *mcr-1-9*, whereas *arnT* encodes a 4-amino-4-deoxy-L-arabinosyltransferase capable of modifying LPS with L-aminoarabinose (L-Ara4N) also lowering the negative charge of LPS (23). These LPS modifications are universally described to provide resistance towards amphipathic cationic peptides. Other modifications implicated in AMP resistance include palmitoylation of Lipid A and removal of phosphates from LPS (24-26). Finally, colistin resistance through complete loss of LPS in *E. coli* and *A. baumannii* has been described (27, 28).

Here we provide evidence that a <u>Ch</u>arge change <u>I</u>ndependent <u>P</u>eptide (ChIP; formerly known as BP214 (29)) has broad-spectrum activity against multiple species of MDR Gram-negative pathogens. ChIP is a linear cationic amphipathic AMP consisting of 11 D-amino acids with the sequence kklfkkilryl-NH_2_ (29). ChIP has a molecular weight (MW) of 1449.89 g/mol and a net positive charge of 6 (at pH 7) comparable to colistin with a MW of 1155.45 g/mol and a net positive charge of 5. We provide evidence that ChIP indeed targets the Gram-negative envelope and that this must be independent of charge due to its activity against colistin resistant strains. Because ChIP is comprised of D-amino acids, protease digestion is not expected to reduce bioavailability. Data indicate that protein binding does not inhibit ChIP activity, but high ChIP concentrations have toxic effects on cultured HeLa cells. However, we demonstrate that a therapeutic window exist that can be exploited in a clinical setting, providing evidence that ChIP is highly efficient against *E. coli* and *A. baumannii* in an intraperitoneal (IP) mouse infection model.

## Results

### ChIP is active against multiple ESKAPE pathogens

We determined the minimum inhibitory concentration (MIC) of ChIP against a small panel of clinical and reference/laboratory isolates (Table 1). ChIP was active against ATCC reference strains of *E. coli, K. pneumoniae* and *A. baumannii* with MIC values in the range of 4-8 µg/ml. The MDR clinical *E. coli* strains ST131 (CTX-M-15), AMA817 (NDM-1) and the *mcr-1* carrying ESBL20150072 (30) remained as sensitive to ChIP as the *E. coli* reference ATCC 25922. The same was observed for *A. baumannii* strains resistant to colistin (CR17, RC64, AB167R and AB176R) which were similar to or more sensitive to ChIP than ATCC 19606 as previously described (29). A single clinical isolate of *A. baumannii* (AB176) had a slightly elevated MIC of 16 µg/ml. For *K. pneumoniae* ATCC 700603 (SHV-18) which is MDR but colistin sensitive the ChIP MIC was 8 µg/ml, while the clinical MDR and colistin resistant *K. pneumoniae* ST512 (Δ*mgrB*) strain had a ChIP MIC of 16 µg/ml. Capsule production has previously been implicated in elevated MIC towards antimicrobial peptides (31). This phenotype was confirmed in the two *K. pneumoniae* strains ATCC 700603 and ST512 (S1).

**Table 1.**
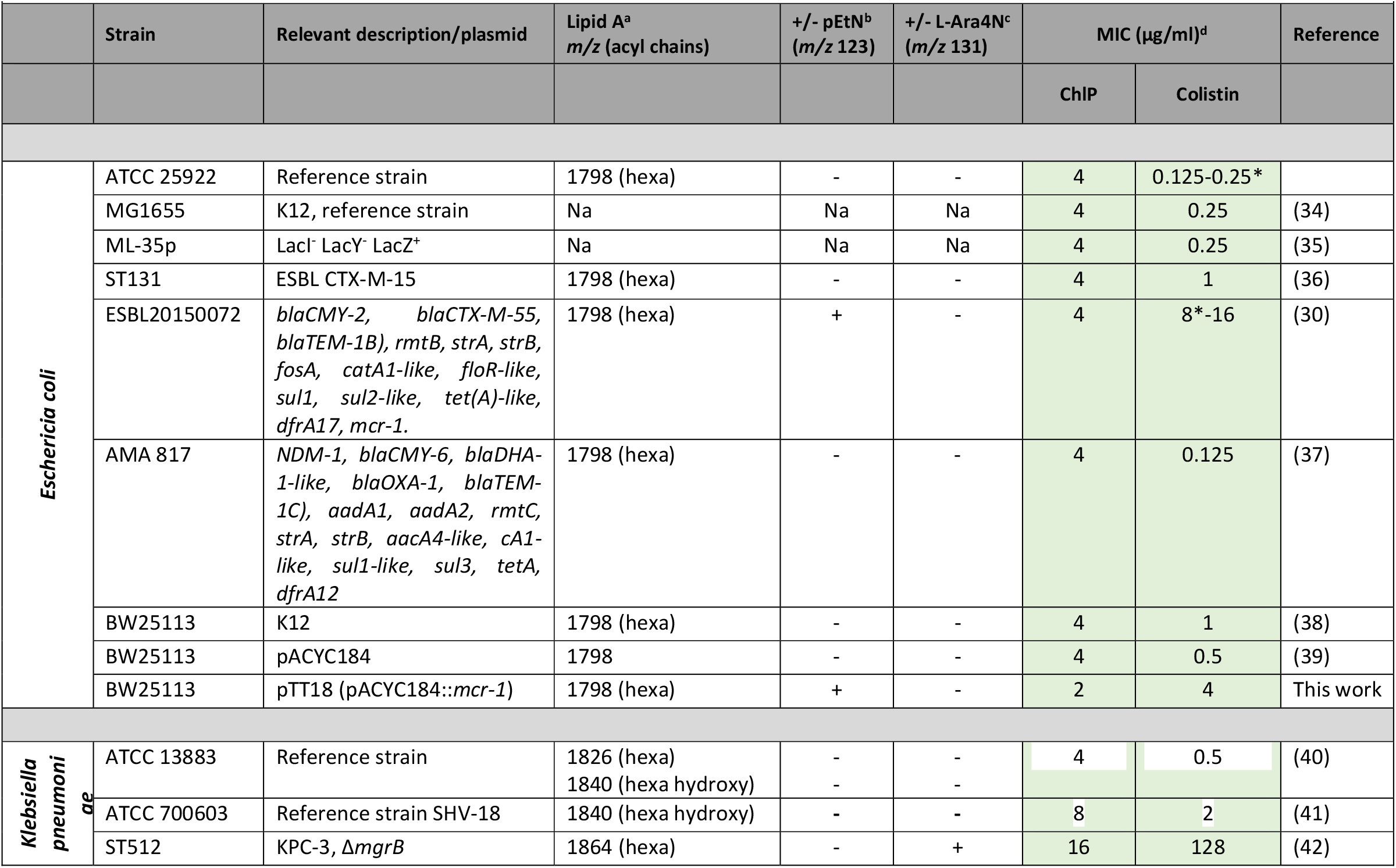
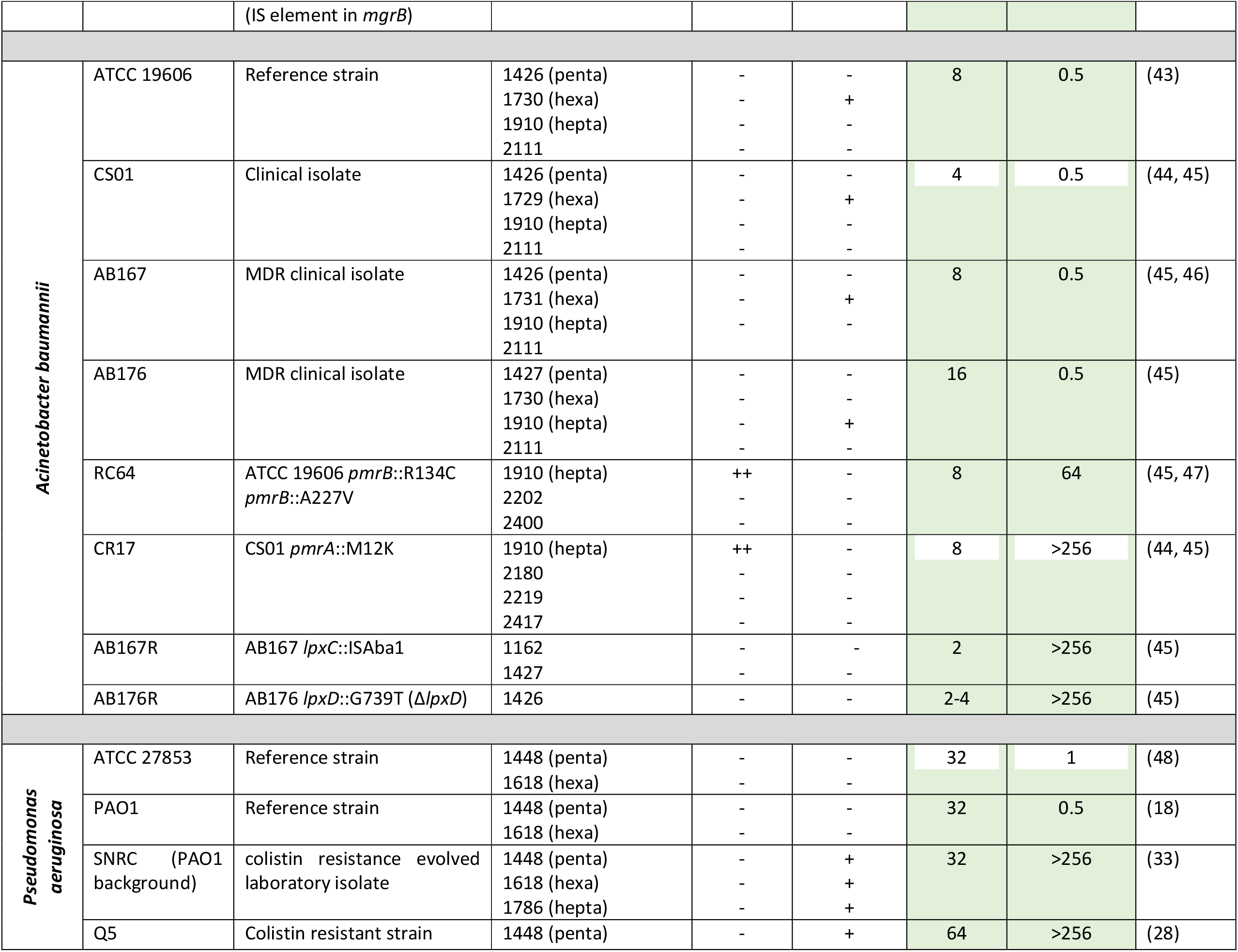
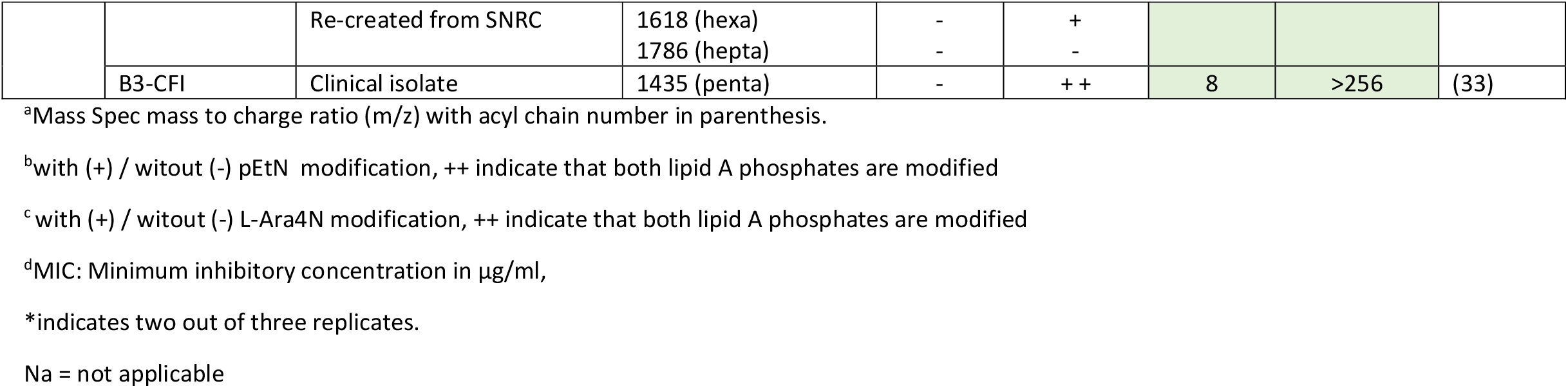
Bacterial species and strain used in this study.

To examine the relationship between colistin resistance and ChIP activity, we cloned the *mcr-1* gene from the clinical strain ESBL20150072 (30) into plasmid pACYC184 resulting in plasmid pTT18. When the *E. coli* strain BW25113 carried pTT18, the colistin MIC increased from 1 µg/ml to 4 µg/ml. This is twofold above the EUCAST defined clinical breakpoint of 2 µg/ml, and result from pEtN modification on Lipid A. The MIC for ChIP was unchanged in the presence of pEtN modification (2-4 µg/ml). The presence of pTT18 in BW25113 resulted in a marked fitness cost as measured by growth rate (62 min) compared to BW25113 (40 min). This is in accordance with previous findings that carriage and expression of the *mcr-1* gene product is a burden to the cell (32).

For *P. aeruginosa* the situation was somewhat different as this background is intrinsically less sensitive to ChIP. Reference strains ATCC 27853 and PAO1 both had a ChIP MIC of 32 µg/ml. Two colistin resistant *P aeruginosa* strains SNRC and Q5 had a ChIP MIC of 32 and 64 µg/ml respectively (33). The SNRC strain is a laboratory evolved colistin resistant strain that carries numerous mutations in genes including; *pmrB, lpxC, opr86, PA5005* and *PA5194* (33). The here named Q5 strain (originally named ABCDE) is PAO1 containing only the 5 recreated mutations in *pmrB, lpxC, opr86, PA5005, PA5194* from SNRC (28). The clinical strain B3-CFI (colistin MIC >256 µg/ml) had a ChIP MIC of 8 µg/ml, implying an inverse relationship between colistin resistance (>256) and ChIP activity in this strain.

Overall, ChIP activity against colistin resistant cells was similar to or lower than against colistin sensitive cells.

### ChIP rapidly kills exponentially growing cells, is stable in human serum, and shows toxicity at high concentrations

Time kill experiments were used to evaluate *in vitro* efficacy of ChIP against exponentially growing *E. coli* (Fig 1). For ATCC 25922 (Fig 1A) and ESBL20150072 (*mcr-1*) (Fig 1B) treatment with 1xMIC ChIP caused a 3-4 log reduction in viable cells followed by a plateau, but with regrowth evident after 3 hours. Treatment with 3xMIC ChlP, reduced cell viability below the detection limit (50 CFU) over the first 5 hours. For comparison, colistin treatment of ATCC 25922 at 1xMIC and 3xMIC resulted in a reduction in CFU to the detection limit. The presence of human male AB serum did not change the MIC of ChIP against *E. coli* ATCC 25922 (Fig 1C). Although this is a crude measure of protein binding/interference, the result clearly shows that ChIP remains active in serum and therefore presumably bind serum proteins poorly.

**Fig 1.**
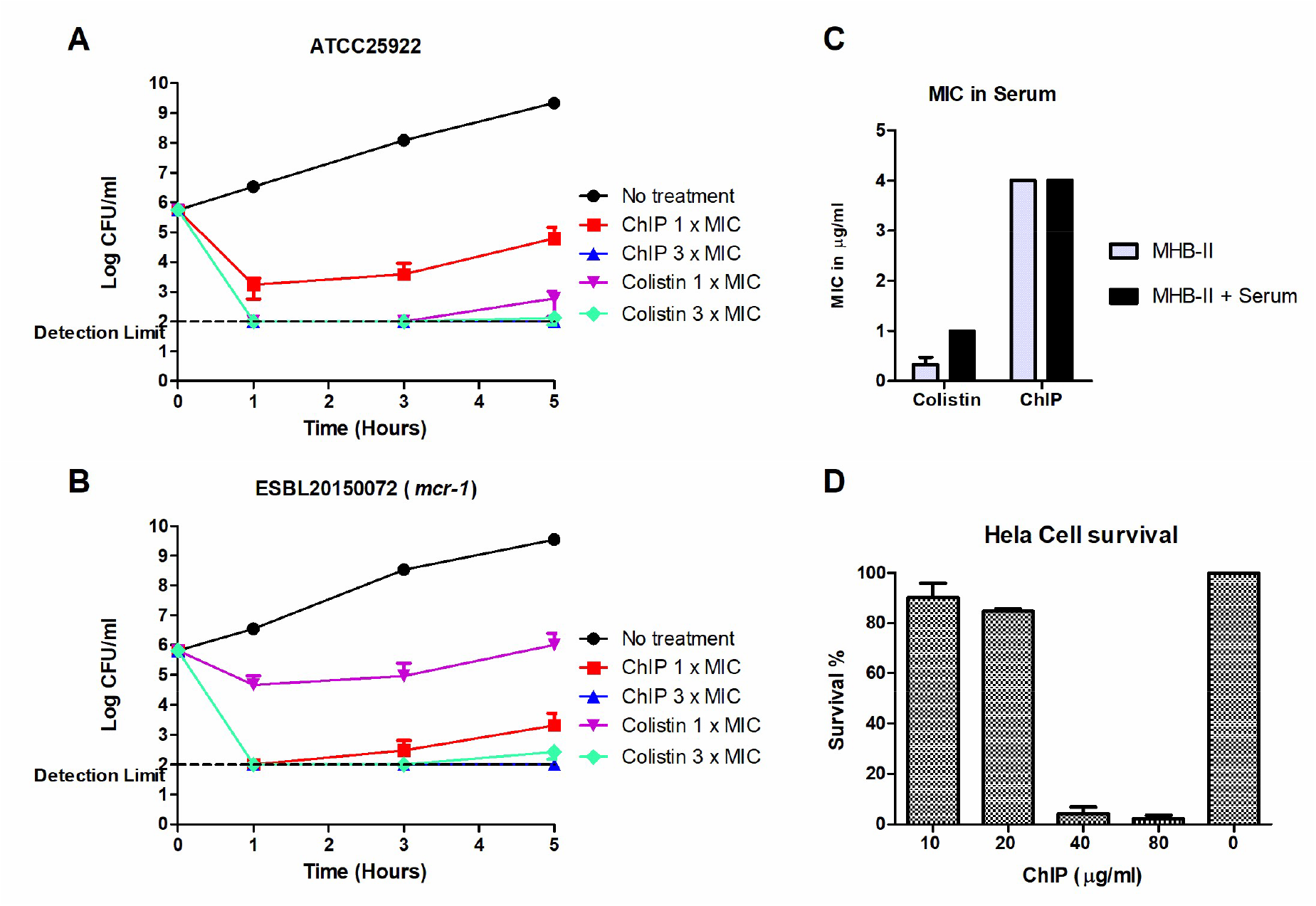
*In vitro* activity and toxicity of ChIP. Treatment of *E. coli* ATCC 25922 (A) and ESBL20150072 (B). Exponentially grown cells are treated with colistin (Col) or ChIP at 1xMIC or 3xMIC at cell densities comparable to MIC (5×10^5^-1×10^6^ CFU/ml). MIC of ChIP and colistin against ATCC 25922 in MHB-II and MHB-II + Human serum (50:50; C). HeLa cell survival after incubation with ChIP at the following concentrations: 10, 20, 40 and 80 µg/ml (D). Experiments were performed in triplicates and the average survival relative to the control is plotted.

ChIP is known to have low hemolytic activity (29), and here we tested toxicity of ChIP on HeLa cell survival following 24 hours of incubation (Fig 1D). More than 80% of HeLa cells survived at ChIP concentrations up to 20 µg/ml, whereas concentrations of 40 µg/ml and above greatly reduced survival. Because the MIC levels for ChIP is significantly lower than 20 µg/ml for most bacterial species and because the molecule has low hemolytic activity, we conclude that a therapeutic window exist for administering this peptide in a clinical setting.

### ChIP is efficient in an *in vivo* infection model

We proceeded to test the antibacterial activity of ChIP in the sepsis/peritonitis mouse infection model. Mice were infected intraperitoneal (IP) with either *E. coli* ATCC 25922 (Fig 2A) or ESBL20150072 (Fig 2B). ChIP treatment (5 mg/kg) reduced peritoneal bacterial titers 2 logs for ATCC 25922 to 1-5×10^5^ CFU/ml, which is similar to treatment with 10 mg/ml colistin. When the ChIP dosage was raised to 10 mg/kg, CFUs were reduced 4 logs to 1×10^3^ CFU/ml. Blood stream CFU for both ChIP treatment groups showed similar reductions (Fig 2A right). Therefore, with treatment at the same concentration, ChIP is superior to colistin in reducing peritoneal bacterial titers.

**Fig 2.**
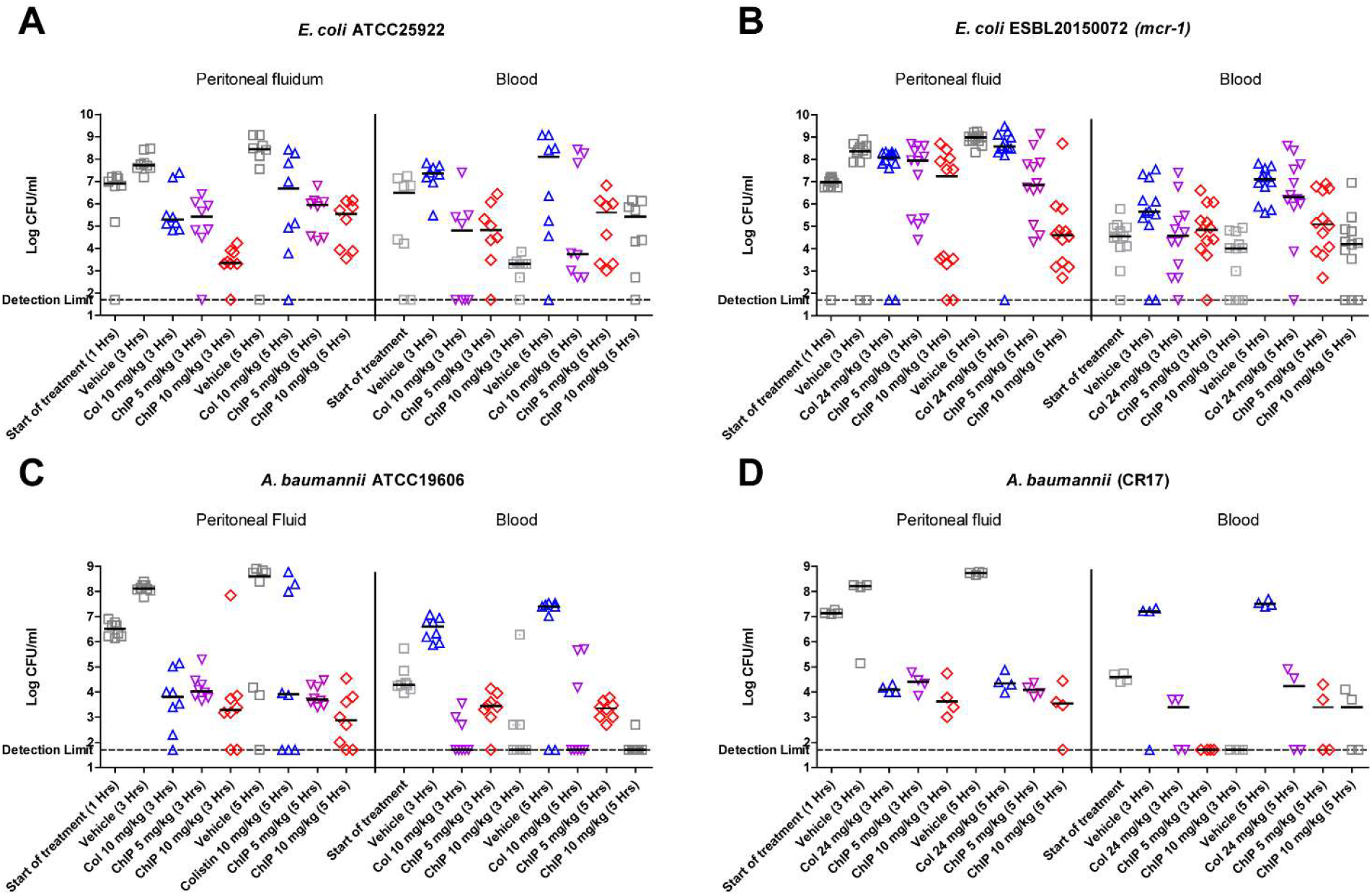
*In vivo* efficacy of ChIP. ChIP efficacy was tested on two strains of the important Gram-negative pathogens *E. coli* and *A. baumannii*. Mice were infected intra peritoneal (IP) and treated with drugs at 1 and 3 hours post infection. Samples were taken at 1 h (start of treatment) and at 3 h and 5 h post infection to determine bacterial titers IP and in the blood stream. Mice infected IP with reference strains *E. coli* ATCC 25922 (A) or *A. baumannii* ATCC 19606 (C) were treated IP with 10 mg/kg colistin (Col) or ChIP (5 or 10 mg/kg). Mice infected with either *E. coli* ESBL20150072 (*mcr-1*; B) or *A. baumannii* CR17 (D), treated with colistin (24 mg/kg) or ChIP (5 or 10 mg/kg). ATCC 25922 and 19606 were tested in duplicate with 4 mice per time point, *E. coli* ESBL20150072 was tested in triplicate (12 mice) per time point whereas CR17 experiments was only carried out as a single experiment. Median CFU/ml (black line) in peritoneum or blood left and right hand side of each graph, respectively.

Against *E. coli* ESBL20150072 (Fig 2B) ChIP treatment (10 mg/kg) likewise resulted in a 3 logs reduction to 1×10^4^ CFU/ml in peritoneal titers 5 hours post infection compared to the vehicle (1×10^8^ CFU/ml) control. Colistin treatment (24 mg/kg) had no effect as expected due to the *mcr-1* genotype. In the ChIP treatment group bacterial blood stream titers remained at 1×10^4-5^ CFU/ml 5 hours post infection, whereas colistin treatment showed increased blood stream titers around ∼1×10^6^ CFU/ml similar to vehicle control.

We proceeded to include two *A. baumannii* isolates for *in vivo* testing, as *A. baumannii* ranks high on the WHO list of critically important pathogens (49). Treatment of ATCC 19606 (Fig 2C) with colistin (10 mg/kg) or ChIP at (5 mg/kg and 10 mg/kg) resulted in bacterial titers around 1×10^3^-1×10^4^ CFU/ml (3-4 log reduction) after 5 hours, relative to the saline control stabilizing around 1×10^8^ CFU/ml after 5 hours. Bacterial blood titers were reduced relative to the vehicle control (Fig 2C right), but with colistin (10 mg/kg) and ChIP (10mg/kg) being able to reduce CFU/ml to the detection limit. The colistin resistant strain CR17 which carries a mutation in the *pmrA* (*basR*) gene, showed similar results to ATCC 19606 both for colistin and ChIP treatment. The reason for colistin treatment being efficient remains puzzling, but we believe it is due to the high colistin concentration being administered (24 mg/ml). For this reason only a single experiment was performed using *A. baumannii* CR17. Overall *in vivo* results clearly demonstrate that ChIP is equal or superior to colistin in reducing the bacterial burden in the mouse peritonitis model.

### ChIP targets and disrupts the Gram-negative bacterial membrane

To narrow down the ChIP target in bacteria, we determined the effect of ChIP on cellular processes by macromolecular incorporation assay. Because these experiments were performed on high cell density cultures (OD_600_ = 0.1, approximately 1×10^8^ CFU/ml) ChIP was tested at 10xMIC. At this concentration, ChIP blocked incorporation of radioactive labelled precursors into DNA, RNA, protein and peptidoglycan of *E. coli* (S2). Because ChIP inhibits all cellular biosynthesis pathways and is bactericidal (Fig. 1), we suspected that the bacterial envelope/membrane integrity is breached, and cells lyse. We therefore proceeded to test the ability of ChIP to permeabilize the inner membrane (Fig 3). We took advantage of the inability of the β-galactosidase substrate ortho-nitrophenyl-β-galactoside (ONPG) to cross the inner membrane of *E. coli*. Consequently, no β-galactosidase activity was detected when adding ONPG to ML-35p cells expressing β-galactosidase at a constitutive high level. However, when cells were treated with ChIP the β-galactosidase activity increased in a dose dependent manner, and to a level higher than that obtained by toluene or colistin treatment (2 µg/ml) (Fig 3B). This indicates that ChIP is more effective than colistin and toluene at permeabilizing *E. coli* cells. ChIP induced permeability for ONPG was accompanied by loss of viability, but without lysis within the first 30 min of treatment (Fig 4).

**Fig 3.**
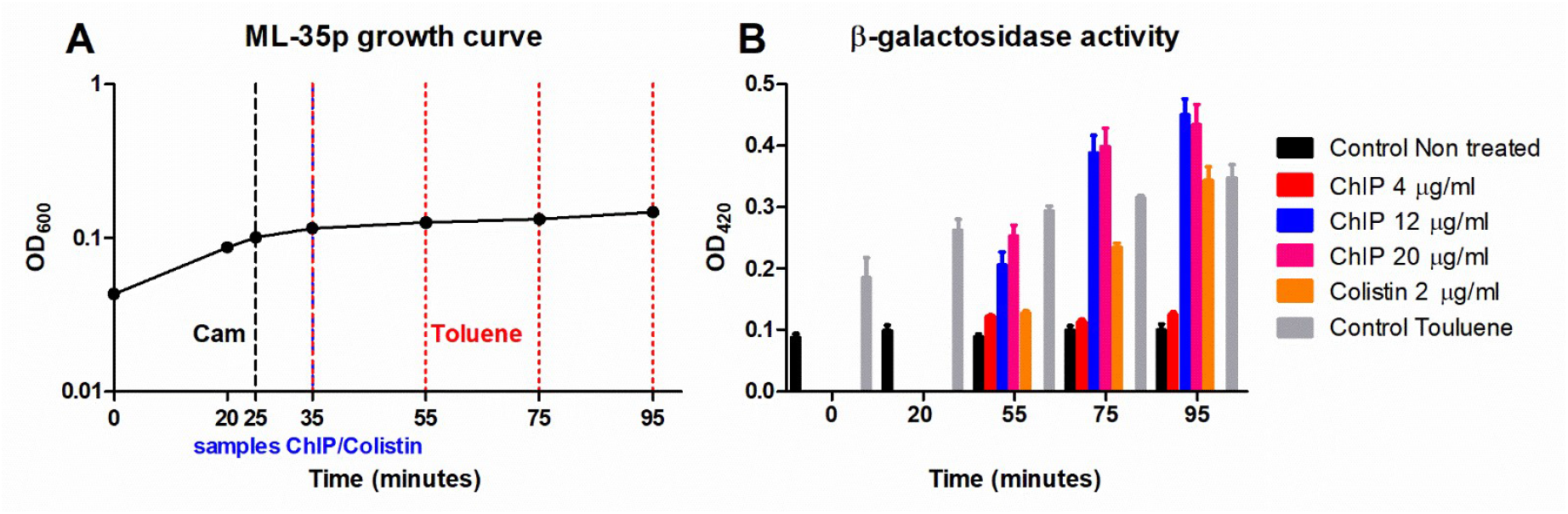
Permeabilization of *E. coli* by ChIP. **A**: The mean growth rate of triplicate cultures of *E. coli* ML-35p (LacI^-^ LacY^-^ LacZ^+^) grown in MHB-II. At time point 25 minutes (black dotted line) the cultures are treated with the bacteriostatic drug chloramphenicol (Cam) 200 µg/ml to block further protein synthesis. 10 minutes post cam treatment (35 minutes) (Blue dotted line), 12 samples are taken from each of the three cultures and treated for 20, 40 or 60 minutes with 2 µg/ml of colistin or 4, 12 and 20 µg/ml ChIP. Samples for toluene treatment are taken as controls at time 0, 20, 55, 75 and 95 minutes (red dotted lines). **B**: The β-galactosidase activity of the treatment groups after 15 min incubation in Z-buffer with ONPG 1 mg/ml at room temperature.

**Fig 4.**
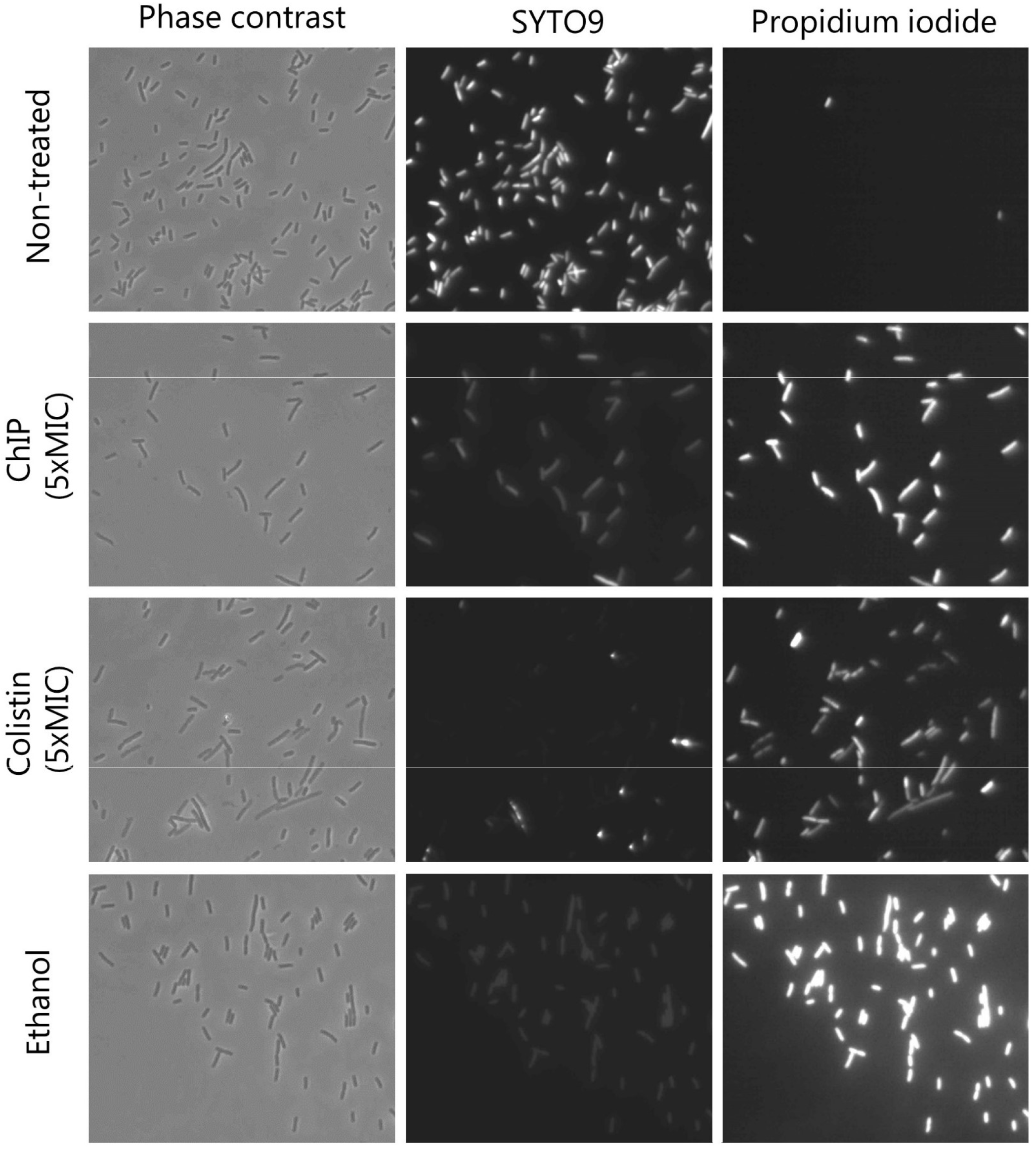
CHIP kill bacterial cell without lysis. ATCC 25922 cells were treated with ChIP, colistin or ethanol and visualized by phase contrast microscopy. Live dead staining performed with SYTO9 - or Propidium iodine and visualized with fluorescence microscopy 30 minutes post treatment as indicated. The pictures are representative of duplicates.

### ChIP activity correlates with the Lipid A acylation pattern

Having established ChIP as a membrane targeting compound, we wanted to explore whether the differences in MIC between the different Gram-negative species can be related to differences in the outer membrane, and proceeded to purify and analyze Lipid A structures using MALDI-TOF MS (Table 1). Across the Gram-negative species and strains analyzed (n = 23), 12 strains carry either pEtN or L-Ara4N modification. All *E. coli* carrying pEtN modification (ESBL20150072 and BW25113::pTT18) remain as sensitive to ChIP as WT (4 µg/ml). Only a single L-Ara4N modified *A. baumannii* (AB176) out of six strains with either pEtN or L-Ara4N modifications had elevated MIC (16 µg/ml) compared to the ATCC 19606 reference strain (8 µg/ml) (table 1). Out of three *P. aeruginosa* strains carrying L-Ara4N modified Lipid A (B3-CFI, SNRC and Q5), only the Q5 strain had a higher MIC (64µg/ml) compared to the ATCC 27853 reference strain (32 µg/ml). While the *P. aeruginosa* (B3-CFI) is in fact more sensitive to ChIP (MIC = 8 µg/ml), than the ATCC 27853 reference strain. One *K. pneumoniae* (ST512) with L-Ara4N modification had a higher MIC (16 µg/ml) compared to ATCC 13883 (4 µg/ml), but it also produces capsule. For these reasons pEtN and L-Ara4N modification were ruled out as drivers of resistance to ChIP. Because lipid A varies slightly among Gram-negative species, depending amongst others on the acyl chain numbers and length of such (50), we proceeded to compare the acyl chain distribution on glucosamine 1 (GlcN1) and glucosamine 2 (GlcN2) of Lipid A from the various strains (Fig 5).

**Fig 5.**
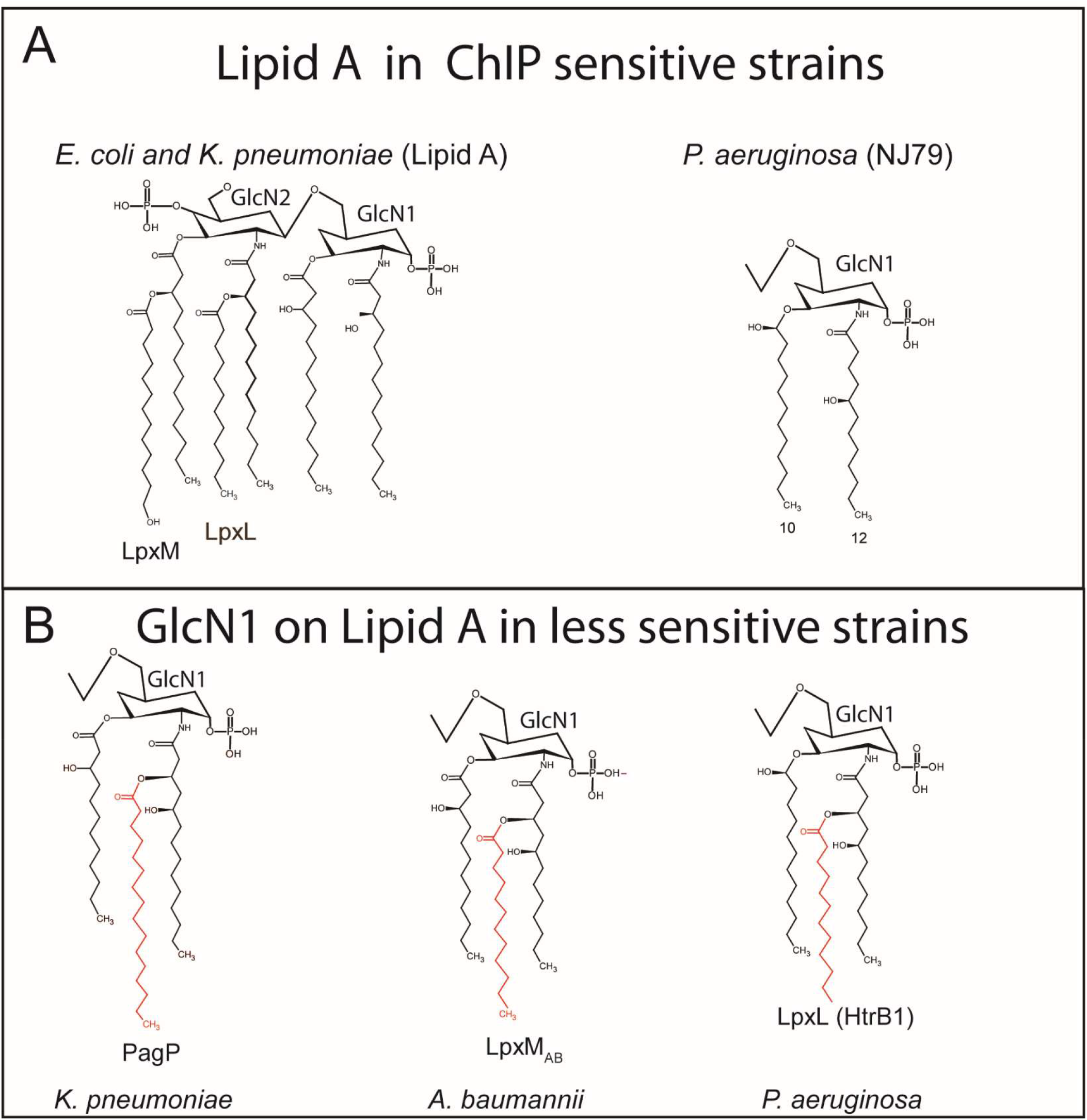
Lipid A structure in relation to lowered ChIP sensitivity. Lipid A structure of Gram-negative cells that are sensitive to ChIP, with only glucosamine 1 (GlcN1) of *P. aeruginosa* shown (A). Acyl chain distribution from four gram-negative species, when these are tolerant/resistant to ChIP (B). Below the different acyl chains are shown the names of proteins responsible for the modifications in the different species.

Most *A. baumannii* strains (ATCC 19606, CS01, AB167 and AB176) primarily carry penta, hexa and hepta acetylated Lipid A as previously published (51, 52) and have MIC values towards ChIP of 4-8 µg/ml (Table 1). The colistin resistant RC64 strain has a highly changed Lipid A profile with *m/z* of 1910 (hepta), *m/z* 2219 and *m/z* 2400, of which the last two to the best of our knowledge, remain undescribed. Nevertheless, cells of all the above strains carried Lipid A species with tri-acylated GlcNI (Fig. 5). Strains AB167R and AB176R were more sensitive to ChIP (2-4 µg/ml) and are LPS deficient due to their *ΔLpxC* and *ΔLpxD* genotypes. This suggests that LPS of the outer membrane constitutes a barrier for ChIP.

*P. aeruginosa* is known to mainly carry 10-12 carbon penta (*m/z* 1448) or hexa (*m/z* 1618) acetylated Lipid A species (53, 54), corresponding to GlcN2 with three acyl chains and GlcN1 with two or three acyl chains (Fig. 5AB). Among the strains analyzed in this study this was the case for ATCC 27853 and PA01. For the colistin resistant strains SNRC and Q5, an additional lipid A species of m/z 1786 was found, corresponding to a hepta acetylated form, with four acyl chains on GlcN1 (53). All of these strains were resistant to ChIP with MIC values of 32-64 µg/ml. Interestingly, the colistin resistant B3-CFI strain, presented Lipid A with a mass solely corresponding to penta acetylated Lipid A (*m/z* 1435 in an L-Ara4N modified version), but with only two acyl chains on GlcN1 (Fig. 5A). This strain is sensitive to ChIP with a MIC of 8 µg/ml indicating the presence of 3 or more acyl chains on GlcN1 drive ChIP resistance. This observation was corroborated by looking at the tested *K. pneumonia* and *E. coli* strains. The majority of these strains carry hexa acetylated Lipid A, with two acyl chains on GlcN1 and four on GlcN2 (Fig. 5), but with a different m*/z* size due to a difference in ratio distribution of carbon chain length between *E. coli* and *K. pneumoniae* (55) (Table 1). These strains were all sensitive to ChIP with MIC values of 2-4 µg/ml (Table 1). The exception was *K. pneumonia* strain ST512 (KPC-3) that had a 4-fold increase in MIC relative to the reference strain ATCC 13883 (Table 1). ST512 carries an alternative hexa acetylated Lipid A (*m/z* 1866) with 3 acyl chains on both GlcN1 and GlcN2 as previously reported (55).

Taken together these data indicate LPS of the outer membrane as an obstacle to ChIP’s ability to kill bacteria. Strains lacking LPS in the outer membrane displayed increased sensitivity to ChIP, while strains with increased MIC values towards ChIP, carry Lipid A with differing acyl chain patterns specified by a more densely acylated GlcN1. We observed no correlation between pEtN or L-Ara4N modification and sensitivity to ChIP.

## Discussion

In the present work, we show that the efficacy of the cationic antimicrobial peptide ChIP is independent of the surface charge change imposed by pEtN and L-Ara4N modification to Lipid A as ChIP remains active against multiple Gram-negative species resistant to colistin. ChIP seemingly kills bacterial cells by permeabilizing the plasma membrane. Data indicate that distribution and level of acetylation on Lipid A determine ChIP sensitivity. Importantly ChIP is active in an *in vivo* infections model against clinical colistin resistant *E. coli* and *A. baumannii*.

### Encapsulation as protection towards ChIP

It is well established that many clinical strains of *K. pneumoniae* (56) produce capsules. Indeed, the mucoid *K. pneumoniae* strains (ATCC 700603 and ST512) had increased MIC values relative to the non-mucoid ATCC 13883 reference strain. Because the mucoid phenotype is associated with capsule formation, we propose that the presence of a capsule provide additional protection against ChIP, similar to what was observed for other AMP (57).

### Surface charge changes imposed by *mcr-1* and or *EptA/ArnT* do not provide protection against ChIP

Loss of LPS in *A. baumannii* resulted in increased sensitivity towards ChIP, indicating that the outer membrane serves as first line of defense against ChIP. At the same time ChIP treatment permeabilizes the inner membrane of *E. coli* and this is likely to be the cause of its bactericidal mode of action. It is therefore conceivable that ChIP acts in a manner similar to colistin with an initial interaction with the LPS layer of the outer membrane that actually constitute an initial barrier followed by ChIP penetration and access to the inner membrane that is permeabilized. We suggest that this permeabilization is the cause of cell death. Lipid A modification via either pEtN addition or L-Ara4N to the 1’ or 4’ positions on Lipid A result in a net charge change to the envelope of Gram-negative pathogens that leads to colistin resistance (20). The same changes to Lipid A did not influence susceptibility to ChIP, and this argues against charge changes as driver of ChIP resistance. Our data seemingly contrasts a recent study (58), which showed high genetic diversity in *E. coli* with evolved resistance to different cationic AMP, but concluded that surface charge changes were the main drivers of resistance. A similar conclusion was reached by another group, working with Arenicin-3 with activity against colistin resistant strains (59). These differences emphasizes the need for a better understanding of cationic antimicrobial peptides’ mechanism of action and resistance development in general.

### Acylation pattern is important for ChIP activity

When we related ChIP activity to LPS acylation pattern, some important observations were made for the individual species. The intrinsically ChIP resistant *P. aeruginosa* strains carry two forms of Lipid A, a penta and a hexa acetylated form (53). The hexa variant is interesting, because it spatially gives *P. aeruginosa* three acyl chains on both GlcN1 and GlcN2. On the other hand, the ChIP sensitive *P. aeruginosa* strain B3-CFI is penta acetylated with only two acyl chains on GlcN1 (Fig. 5). This indicates that the presence of LPS types with three acyl chains on GlcN1 confers resistance to ChIP. Observations from both *A. baumannii* and *K. pneumoniae* support this hypothesis. Most *A. baumannii* strains including ATCC19606 displayed intermediate sensitivity to ChIP and carry both hexa and hepta acetylated Lipid A. Importantly, the hepta form carries a GlcN1 acyl chain distribution similar to ChIP resistant *P. aeruginosa*, which is produced by LpxM during Lipid A synthesis (Fig. 5). Loss of LPS in the two *A. baumannii* strains, AB167R and AB176R, leads to increased sensitivity towards ChIP. Among *K. pneumoniae* strains, ST512 has the highest MIC towards ChiP. This strain carries an alternative hexa acetylated Lipid A, in which the acylation pattern is similar to the hexa form found in *P. aeruginosa* with three acyl chains on GlcN1 (Fig. 5). *K. pneumoniae* has the capability to modify GlcN1 with an extra acyl chain via *pagP* (Lipid A palmitoyltransferase) (24, 55, 60) that utilizes misplaced phospholipids in the OM to modify Lipid A on the fully synthesized LPS (24, 25, 61).

We therefore propose that it is the distribution of acyl chains that drives sensitivity to ChIP and suggest that strains carrying an elevated level of Lipid A types with three acyl chains on GlcN1 become resistant to ChIP (Fig 5). We speculate that the presence of Lipid A carrying tri-acylated GlcN1 allows for a denser outer membrane that is less susceptible for lysis/permeabilization by ChIP. In agreement with this, all tested *E. coli* strains carry mainly hexa acetylated Lipid A with two acyl chains on GlcN1 and all are sensitive to ChIP. Previous studies have also shown differences in relation to the acylation pattern on LPS between Gram-negative strains (24) and palmitoylation has been associated with AMP resistance (21). We do acknowledge that such cross-species comparisons are questionable and this is even the case for different isolates within the same species. Especially clinical isolates pose problems in answering questions related to antimicrobial sensitivity as they differ genetically and may possess different characteristics that interfere with AMP activity. Future studies should therefore focus on isogenic strains and should include measuring the actual concentrations of Lipid A variants to clarify acylation pattern vs. charge changes in relation to activity.

### ChIP efficiently kills colistin resistant *E. coli* and *A. baumannii in vivo*

ChIP efficiently reduced the bacterial burden of *E. coli* and *A. baumannii in vivo* in the mouse peritonitis model, with an activity equal to or better than colistin that is currently the last resort antibiotic against complicated infections caused by Gram-negative bacteria. Recently, a modified version of the amphipathic peptide Arenicin was also found to efficiently reduce the bacterial burden in mice infected with *E. coli* or *P. aeruginosa* (59). While Arenicin-3 was tested in several infection models (peritonitis, pneumonia and urinary tract infection) and showed a 7 log reduction in bacterial burden that lasted for days, our data show a more moderate effect of ChIP, with a bacterial reduction of up to 5 logs intraperitoneal after two-dose treatment. Although, Arenicn-3 (β-hairpin) and ChIP (α-helical) differ structurally, they are both envelope acting against MDR colistin resistant Gram-negative pathogens that have modified LPS with pEtN or L-Ara4N. ChIP has yet to undergo considerable optimization, and could therefore possibly be optimized further for activity or to avoid issues of instability and toxicity. This could be by PEGylating the peptide to increase stability and serum elimination half-life (62) or other modification such as peptidomimetics to lower possible toxicological issues (63). Therefore, further toxicological studies and more clinically relevant *in vivo* studies on a second-generation ChIP compounds are needed. ChIP nonetheless remains highly interesting especially in relation to colistin resistance and *mcr-1* and as such AMP are still considered highly relevant in the search for new antibiotics (64).

## Materials and methods

### Bacterial strains and growth conditions

All bacterial strains and plasmids used in this study are listed in Table 1. Bacteria were grown in cation adjusted Müller-Hinton broth (MHB-II; Sigma-Aldrich) for MIC or Luria-Bertani (LB) broth at 37°C, supplemented with relevant antibiotics when required. For macromolecular incorporation measurements cells were grown in AB minimal medium (65) supplemented with 10 μg/ml thiamine and 0.2% glucose without Cas amino acids. Except for glucosamine incorporation where 0.2% glycerol was used as carbon source. Growth rate of the *mcr-1* carrying BW25113 strain was determined in MHB-II on exponentially growing cells by measuring OD_600_ over time.

### Macromolecular incorporation

Macromolecular synthesis was determined by incorporation of 3H-uridine, 3H-thymidine, 3H-arginine or 3H-glucosamine into DNA, RNA, proteins and cell wall respectively, as described previously (66). Briefly, 500 µl samples of exponentially growing MG1655 cells were incubated with 0.375 µCi of the relevant precursor for 2 minutes (uridine), 4 minutes (thymidine and arginine), and 20 minutes (glucosamine). Reactions were stopped by adding 5 ml of ice-cold 5% TCA containing 0.1 M NaCl. Samples were filtered (ADVANTEC®, 25mm, lot no: 80302708) washed twice and dried overnight in scintillation vials (PV1AS, MERIDIAN, BIOTECHNOLOGIES Ltd) before counting on a HINDEX 300SL in 5 ml of scintillation fluid (Scintillation fluid, ULTIMA GOLD^™^, Perkin Elmer, Inc. LOT No.: 77-18151).

### Permeabilization assay

Exponentially growing cells of *E. coli* ML35 (LacI^-^ LacY^-^ LacZ^+^) were treated with chloramphenicol (200 µg/ml) to stop protein synthesis. Chloramphenicol treated samples were incubated with ChIP (4, 12 and 20 µg/ml) and colistin (2 µg/ml) for 20, 40 or 60 minutes (12 samples total). 200 µL peptide treated cells were added to 1 ml Z-buffer (0.06M Na_2_HPO_4_, 0.04M NaH_2_PO_4_, 0.01M KCL, 0.001M MgSO_4_ and pH = 7) containing 1 mg/ml Ortho-Nitrophenyl-β-galactoside (ONPG). Samples were left at room temperature 15 minutes before stopping color development by addition of 0.5 ml Na_2_CO_3_ (1M). Toluene treated cells were included as a control for permeabilized cells. β-galactosidase activity was detected by OD_420_ measurements.

### Live/Dead staining

Exponentially growing *E. coli* ATCC25922 were treated for 30 min with either colistin (5xMIC) or ChIP (5xMIC) at OD_600_ = 0.1. An untreated culture was used as a control. 20 ml of treated or untreated culture was concentrated by centrifugation at 5,000 × g for 10 minutes, washed and re-suspended in 10 ml 0.9% NaCl. All samples were incubated at room temperature for 1 hour, mixing every 15 minutes, before washing twice with 10 ml 0.9% NaCl. After washing, the samples were re-suspended in 5 ml 0.9% NaCl, except for ChIP treated samples that were suspended in 1 ml 0.9% NaCl. Sample staining was performed according to the manufacturers instruction (LIVE/DEAD(tm) *Bac*Light(tm) Bacterial Viability Kit, for microscopy & quantitative assays), and incubated for 15 minutes in the dark. 5 μl of the bacterial suspension was placed on an agar pad and analyzed by fluorescence microscopy using an AxioImager Z1 microscope (Carl Zeiss MicroImaging, Inc), with a × 100 objective and a Hamamatsu ORCA-ER C4742-80-12AG camera.

### Construction of *mcr-1* carrying isogenic *E. coli* of non-clinical origin

The *mcr-1* gene was amplified from the clinical isolate ESBL20150072 (European Nucleotide Archive, accession number; SAMEA104060671) using primers: 5’-TTTTGGATCCCGGTTTTCGGGCTTTTTA-3’ and 5’-ATATGTCGACTCAGCGGATGAATGCGGT-3’ and Phusion polymerase (Thermo Fisher Scientific). PCR product was digested with *Bam*HI and *Sal*I and ligated into pACYC184 digested with the same enzymes (Thermo Fisher Scientific).

### MALDI-TOF Mass-spec verification of Lipid A modification

Membrane lipids were purified as previously described (67). Briefly, 20 ml of exponentially growing culture (MHB-II media) was spun down (12,000 g 10 min) and re-suspended in 400 µl 100 mM sodium acetate buffer pH 4, incubated for 30 min at 100°C (vortexing every 5-10 min) and cooled to room temperature on ice. The supernatant was discarded after centrifugation at 8,000 g for 5 min. The pellet was washed in 1 ml 96% ethanol and insoluble lipids were extracted with 100 µl chloroform/methanol/water (12:6:1, v/v/v). Following centrifugation (5,000 g for 5 min) the supernatant was spotted onto MALDI-TOF MS target plate using 2,5-dihydroxybenzoic acid as matrix and samples were run on a Bruker autoflex mass spectrometre and analysed using Flexanalysis Bruker.

### Minimum inhibitory concentrations determination

The minimum inhibitory concentration (MIC) was determined as previously described (68), with minor modifications. Briefly, exponentially growing cultures in MHB-II (37°C) were diluted to an OD_600_ = 0.002, approximately 1×10^6^ colony forming units (CFU) per ml. 100 µl of bacterial suspension was added to polypropylene microtiter plates containing 100 µl of Colistin or ChIP dilutions. Giving a final bacterial inoculum of ∼ 5×10^5^ CFU/ml. Colistin sulfate salt (Sigma-Aldrich) and ChIP (Genscript >95% purity) was dissolved in H_2_O. ChIP MIC in serum was determined in the presence of 50% (heat inactivated) human serum (Sigma) and 50% MHB-II. All experiments were performed in triplicate.

### Time kill experiments

Exponentially growing ATCC 25922 and ESBL20150072 were diluted in fresh MHB-II (37°C) to OD_600_ = 0.001 approximately 5×10^5^ CFU/ml (60 ml). Cultures were split into individual flasks and treated with peptides: Colistin (Col) or ChIP at 1xMIC and 3xMIC. At times 0, 1, 3 and 5 hours, 250 µl culture was removed, washed twice in 0.9% saline, before resuspension in 250 µl saline. A 10-fold dilution series was made in 0.9% saline and 3×10 µl of each dilution was spotted onto LB agar without selection. After 24 hours at 37°C CFU were counted.

### Phenotypic characterization of capsule formation

Capsule formation was assessed by streaking on solid lactose agar plates (‘Bromthymole Blue plates’ based on a modified Conradi-Drigalski medium containing 10 g/L detergent, 1 g/L Na_2_S_2_O_3_·H_2_O, 0.1 g/L bromothymol blue, 9 g/L lactose and 0.4 g/L glucose, pH 8.0; Statens Serum Institut, Copenhagen, Denmark).

### Antimicrobial peptide toxicity to HeLa cells

HeLa cells were cultured in Dulbecco’s Modified Eagles Medium (DMEM) (Sigma) supplemented with 10% fetal calf serum (Sigma) and 2 mM glutamine at 37°C in a humid incubator with 5% CO_2_. Cell survival after 24 hours with various amounts of ChIP in DMEM without serum was determined by staining with Trypan Blue Stain (Gibco/Life Technologies). Experiments were performed in triplicate.

### *In vivo* sepsis/peritonitis mouse infection model

Outbred female NMRI mice (NMRI; weight 26–30 g; Taconic, Denmark) were used throughout the study, as previously described (69, 70). We decided to dose ChIP in IP, comparable to colistin intravenous dosages. Colistin is recommended at a dose of 5-10 mg/kg, according to the European Medicines Agency (EMA) (71). The animals were kept in Macrolon type III cages in groups of six and allowed free access to feed and water. Experiments were initiated after an acclimatization period of 7 days. Inoculation (bacterial infection) was performed by intraperitoneal (IP) injection of 0.5 ml bacterial suspension containing 2×10^7^ CFU/ml and 5% (wt/vol) mucin (Sigma-Aldrich). During experiments, groups of eight mice per cage constituting one experimental unit were randomly assigned to treatment (5 mg/kg ChIP and 10 mg/kg ChIP or colistin 10 mg/kg and 24 mg/kg) or no treatment (vehicle). Peptide and colistin was administered as a single bolus injection IP (volume injected 0.2 ml) at 1 hours and again 3. At various times post-infection, ∼200 µL blood from the jaw/cheek was collected in Eppendorf tubes with anti-coagulate (Sarstedt, Nümbrecht, Germany). Peritoneal lavage was performed by injecting 2.0 ml of sterile physiological saline IP. After 1 minute of abdominal massage, the peritoneum was opened and peritoneal lavage fluid (PLF) withdrawn. PLF was immediately placed in an insulated 4^°^C cooling box for transportation and kept on ice at 4^°^C. The mouse peritonitis model was repeated in duplicates for *E. coli* ATCC 25922 and *A. baumannii* ATCC 19606 nad in triplicate for *E. coli* ESBL20150072, while only a single experiment was carried out on *A. baumannii* CR17. Animals were sacrificed at various times prior to and following infection. Data from repeated experiments were pooled for analyses. Quantification of bacterial load was performed by spot plating of blood and peritoneal fluid samples from the mice. Dilution series were made in sterile 0.9% NaCl solution, from which 20 μl was spotted on solid lactose agar plates (Bromthymole Blue plates, SSI Diagnostica). Bacterial counts were recorded after overnight incubation at 37^°^C in ambient air. The detection limit was 50 CFU/ml. The clinical condition of the mice was determined using a self-developed clinical score. For this score, mouse behavior and appearance were observed and evaluated for the parameters piloerection, attitude, locomotion, breathing and curiosity and awarded a score from 0 to 5 by a blinded observer. 0 = no clinical signs, 1 = slightly ruffled fur, 2 = ruffled fur and strenuous breathing, 3 = 2 plus lethargy, 4 = 3 plus hunched posture and decreased activity, and 5 = moribund. No significant difference in the clinical score between groups was observed 2 and 4 h post treatment. All mice were recorded for score 2.

## Ethical statement

All animal experiments were approved by the Danish Animal Experimentation Inspectorate (license no. 2017-15-0201-01274) and performed according to institutional guidelines. The mice regularly observed and scored for signs of distress. Humane endpoints constituted signs of irreversible sickness; mice euthanized upon presentation of any of these signs.

## Funding and support

We would like to thank the following foundations for financial support of this project. The Novo Nordisk Foundation: Novo Nordisk Fonden: NNF16OC0021700 (Anders Løbner-Olesen) and Novo Nordisk Foundation: NNF16OC0023482 (Niels Frimodt-Møller and Anders Folkesson). Furthermore, we would like to thank Kirsten and Freddy Johansens Foundation (Anders Løbner-Olesen) for financial support.

## Author Contributions

**Conceptualization:** Thomas T. Thomsen, Anders Løbner-Olesen

**Data curation:** Henrik Hasman

**Funding Acquisition:** Anders Løbner-Olesen, Niels Frimodt-Møller, Anders Folkesson

**Investigation:** Thomas T. Thomsen, Mette Kolpen, Vinoth Wigneswaran, Ulrik Kromann, Anna E. Ebbensgaard, Anette M. Hammerum, Henrik Hasman, Stine Radmer and Kasper N. Kragh

**Methodology:** Thomas T. Thomsen, Mette Kolpen, Vinoth Wigneswaran, Ulrik Kromann, Anna E. Ebbensgaard, Anette M. Hammerum, Henrik Hasman, Stine Radmer, Kasper N. Kragh, Niels Frimodt-Møller, Anders Folkesson

**Visualization:** Thomas T. Thomsen, Mette Kolpen and Anders Løbner-Olesen

**Writing-Original Draft Preparation:** Thomas T. Thomsen, Anders Løbner-Olesen

**Writing – Review & Editing:** Thomas T. Thomsen, Mette Kolpen, Vinoth Wigneswaran, Ulrik Kromann, Anna E. Ebbensgaard, Anette M. Hammerum, Henrik Hasman, Stine Radmer, Kasper N. Kragh, Rasmus Hartmann-Petersen, Paul Robert Hansen, Anders Folkesson, Niels Frimodt-Møller and Anders Løbner-Olesen.

## Supporting information captions

**S1. Phenotypic capsule determination**. S1A. *K. pneumoniae* strains ATCC 13883, ATCC 700603 and ST512. *K. pneumoniae* ST512 and ATCC 700603 clearly appear mucoid on solid agar compared to the ATCC 13883 strain, this is typical of capsule formation. Fig S1B shows non mucoid E. coli strains for comparison (ATCC 25922, ESBL20150072, CTX-M-15 and NDM-1).

**S2. Macromolecular incorporation assay**. Figures S2A, S2B, S2C and S2D shows the results ChIP treatment on major cellular biosynthesis processes (DNA, RNA, Protein and cell wall biosynthesis) respectively. Y axis shows counts per minute (CPM) divided by OD_450_ (when sample was taken). The x axis shows time in minutes for experiments. The right graphs 1E, 1F, 1G and 1H shows the OD_450_ measurements (Y axis) plotted over time (x axis). Cells were treated (40 min, dotted line) with ChIP or control drugs (Nalidixic acid (Nal), Rifampicin (Rif), Chloramphenicol (Cam) or ampicillin (Amp). Graphs are from representative experiment.

## Notes

### Competing Interest Statement

The authors have declared no competing interest.

## References

1. WHO. Antimicrobial resistance: global report on surveillance 2014. 2014:257.

2. Boucher HW, Talbot GH, Bradley JS, Edwards JE, Gilbert D, Rice LB, et al. Bad bugs, no drugs: no ESKAPE! An update from the Infectious Diseases Society of America. Clinical infectious diseases : an official publication of the Infectious Diseases Society of America. 2009;48(1):1–12.

3. Fjell CD, Hiss JA, Hancock RE, Schneider G. Designing antimicrobial peptides: form follows function. Nature reviews Drug discovery. 2012;11(1):37–51.

4. Hancock RE. Peptide antibiotics. Lancet. 1997;349(9049):418–22.

5. Jenssen H, Hamill P, Hancock RE. Peptide antimicrobial agents. Clinical microbiology reviews. 2006;19(3):491–511.

6. Chen CH, Lu TK. Development and Challenges of Antimicrobial Peptides for Therapeutic Applications. Antibiotics (Basel, Switzerland). 2020;9(1).

7. Mojsoska B, Jenssen H. Peptides and Peptidomimetics for Antimicrobial Drug Design. Pharmaceuticals (Basel). 2015;8(3):366–415.

8. Perron GG, Zasloff M, Bell G. Experimental evolution of resistance to an antimicrobial peptide. Proceedings Biological sciences. 2006;273(1583):251–6.

9. Zasloff M. Antimicrobial peptides of multicellular organisms. Nature. 2002;415(6870):389–95.

10. Silhavy TJ, Kahne D, Walker S. The Bacterial Cell Envelope. Cold Spring Harbor perspectives in biology. 2010;2(5):a000414.

11. Nikaido H. Molecular Basis of Bacterial Outer Membrane Permeability Revisited. Microbiology and Molecular Biology Reviews. 2003;67(4):593–656.

12. Henderson JC, Zimmerman SM, Crofts AA, Boll JM, Kuhns LG, Herrera CM, et al. The Power of Asymmetry: Architecture and Assembly of the Gram-Negative Outer Membrane Lipid Bilayer. Annual review of microbiology. 2016;70:255–78.

13. Falagas ME, Kasiakou SK. Colistin: the revival of polymyxins for the management of multidrug-resistant gram-negative bacterial infections. Clinical infectious diseases : an official publication of the Infectious Diseases Society of America. 2005;40(9):1333–41.

14. Berglund NA, Piggot TJ, Jefferies D, Sessions RB, Bond PJ, Khalid S. Interaction of the antimicrobial peptide polymyxin B1 with both membranes of E. coli: a molecular dynamics study. PLoS computational biology. 2015;11(4):e1004180.

15. Sabnis A, Hagart KLH, Klöckner A, Becce M, Evans LE, Furniss RCD, et al. Colistin kills bacteria by targeting lipopolysaccharide in the cytoplasmic membrane. eLife. 2021;10.

16. Liu Y-Y, Wang Y, Walsh TR, Yi L-X, Zhang R, Spencer J, et al. Emergence of plasmid-mediated colistin resistance mechanism MCR-1 in animals and human beings in China: a microbiological and molecular biological study. The Lancet Infectious Diseases.

17. Davies M, Walsh TR. A colistin crisis in India. Lancet Infect Dis. 2018;18(3):256–7.

18. Ling Z, Yin W, Shen Z, Wang Y, Shen J, Walsh TR. Epidemiology of mobile colistin resistance genes mcr-1 to mcr-9. The Journal of antimicrobial chemotherapy. 2020;75(11):3087–95.

19. Guo L, Lim KB, Gunn JS, Bainbridge B, Darveau RP, Hackett M, et al. Regulation of lipid A modifications by Salmonella typhimurium virulence genes phoP-phoQ. Science (New York, NY). 1997;276(5310):250–3.

20. Raetz CRH, Reynolds CM, Trent MS, Bishop RE. LIPID A MODIFICATION SYSTEMS IN GRAM-NEGATIVE BACTERIA. Annual review of biochemistry. 2007;76:295–329.

21. Simpson BW, Trent MS. Pushing the envelope: LPS modifications and their consequences. Nature reviews Microbiology. 2019;17(7):403–16.

22. Froelich JM, Tran K, Wall D. A pmrA constitutive mutant sensitizes Escherichia coli to deoxycholic acid. Journal of bacteriology. 2006;188(3):1180–3.

23. Trent MS, Ribeiro AA, Doerrler WT, Lin S, Cotter RJ, Raetz CR. Accumulation of a polyisoprene-linked amino sugar in polymyxin-resistant Salmonella typhimurium and Escherichia coli: structural characterization and transfer to lipid A in the periplasm. The Journal of biological chemistry. 2001;276(46):43132–44.

24. Bishop RE, Gibbons HS, Guina T, Trent MS, Miller SI, Raetz CR. Transfer of palmitate from phospholipids to lipid A in outer membranes of gram-negative bacteria. The EMBO journal. 2000;19(19):5071–80.

25. Boll JM, Tucker AT, Klein DR, Beltran AM, Brodbelt JS, Davies BW, et al. Reinforcing Lipid A Acylation on the Cell Surface of Acinetobacter baumannii Promotes Cationic Antimicrobial Peptide Resistance and Desiccation Survival. mBio. 2015;6(3):e00478.

26. Needham BD, Trent MS. Fortifying the barrier: the impact of lipid A remodelling on bacterial pathogenesis. Nature reviews Microbiology. 2013;11(7):467–81.

27. Moffatt JH, Harper M, Harrison P, Hale JD, Vinogradov E, Seemann T, et al. Colistin resistance in Acinetobacter baumannii is mediated by complete loss of lipopolysaccharide production. Antimicrobial agents and chemotherapy. 2010;54(12):4971–7.

28. Moosavian M, Emam N, Pletzer D, Savari M. Rough-type and loss of the LPS due to lpx genes deletions are associated with colistin resistance in multidrug-resistant clinical Escherichia coli isolates not harbouring mcr genes. PloS one. 2020;15(5):e0233518.

29. Alberto Oddo TT, Susanne Kjelstrup, Ciara Gorey, Henrik Franzyk, Niels Frimodt-Møller, Anders Løbner-Olesen, and Paul Hansen. An all-D amphipathic undecapeptide shows promising activity against colistin-resistant strains of Acinetobacter baumannii and a dual mode of action. AAC01966-15R1. 2015.

30. Hasman H, Hammerum AM, Hansen F, Hendriksen RS, Olesen B, Agerso Y, et al. Detection of mcr-1 encoding plasmid-mediated colistin-resistant Escherichia coli isolates from human bloodstream infection and imported chicken meat, Denmark 2015. Euro surveillance : bulletin Europeen sur les maladies transmissibles = European communicable disease bulletin. 2015;20(49).

31. Campos MA, Vargas MA, Regueiro V, Llompart CM, Alberti S, Bengoechea JA. Capsule polysaccharide mediates bacterial resistance to antimicrobial peptides. Infection and immunity. 2004;72(12):7107–14.

32. Yang Q, Li M, Spiller OB, Andrey DO, Hinchliffe P, Li H, et al. Balancing mcr-1 expression and bacterial survival is a delicate equilibrium between essential cellular defence mechanisms. Nature communications. 2017;8(1):2054.

33. Jochumsen N, Marvig RL, Damkiaer S, Jensen RL, Paulander W, Molin S, et al. The evolution of antimicrobial peptide resistance in Pseudomonas aeruginosa is shaped by strong epistatic interactions. Nature communications. 2016;7:13002.

34. Guyer MS, Reed RR, Steitz JA, Low KB. Identification of a sex-factor-affinity site in E. coli as gamma delta. Cold Spring Harb Symp Quant Biol. 1981;45 Pt 1:135–40.

35. Eriksson M, Nielsen PE, Good L. Cell permeabilization and uptake of antisense peptide-peptide nucleic acid (PNA) into Escherichia coli. The Journal of biological chemistry. 2002;277(9):7144–7.

36. Cerquetti M, Giufrè M, García-Fernández A, Accogli M, Fortini D, Luzzi I, et al. Ciprofloxacin-resistant, CTX-M-15-producing Escherichia coli ST131 clone in extraintestinal infections in Italy. Clinical microbiology and infection : the official publication of the European Society of Clinical Microbiology and Infectious Diseases. 2010;16(10):1555–8.

37. Hammerum AM, Hansen F, Nielsen HL, Jakobsen L, Stegger M, Andersen PS, et al. Use of WGS data for investigation of a long-term NDM-1-producing Citrobacter freundii outbreak and secondary in vivo spread of blaNDM-1 to Escherichia coli, Klebsiella pneumoniae and Klebsiella oxytoca. Journal of Antimicrobial Chemotherapy. 2016;71(11):3117–24.

38. Datsenko KA, Wanner BL. One-step inactivation of chromosomal genes in Escherichia coli K-12 using PCR products. Proceedings of the National Academy of Sciences of the United States of America. 2000;97(12):6640–5.

39. Chang AC, Cohen SN. Construction and characterization of amplifiable multicopy DNA cloning vehicles derived from the P15A cryptic miniplasmid. Journal of bacteriology. 1978;134(3):1141–56.

40. Cowan ST, Steel KJ, Shaw C, Duguid JP. A classification of the Klebsiella group. Journal of general microbiology. 1960;23:601–12.

41. Rasheed JK, Anderson GJ, Yigit H, Queenan AM, Doménech-Sánchez A, Swenson JM, et al. Characterization of the extended-spectrum beta-lactamase reference strain, Klebsiella pneumoniae K6 (ATCC 700603), which produces the novel enzyme SHV-18. Antimicrobial agents and chemotherapy. 2000;44(9):2382–8.

42. Grundmann H, Glasner C, Albiger B, Aanensen DM, Tomlinson CT, Andrasevic AT, et al. Occurrence of carbapenemase-producing Klebsiella pneumoniae and Escherichia coli in the European survey of carbapenemase-producing Enterobacteriaceae (EuSCAPE): a prospective, multinational study. Lancet Infect Dis. 2017;17(2):153–63.

43. Schaub IG, Hauber FD. A Biochemical and Serological Study of a Group of Identical Unidentifiable Gram-negative Bacilli from Human Sources. Journal of bacteriology. 1948;56(4):379–85.

44. López-Rojas R, Jiménez-Mejías ME, Lepe JA, Pachón J. Acinetobacter baumannii Resistant to Colistin Alters Its Antibiotic Resistance Profile: A Case Report From Spain. The Journal of infectious diseases. 2011;204(7):1147–8.

45. Garcia-Quintanilla M, Pulido MR, Moreno-Martinez P, Martin-Pena R, Lopez-Rojas R, Pachon J, et al. Activity of host antimicrobials against multidrug-resistant Acinetobacter baumannii acquiring colistin resistance through loss of lipopolysaccharide. Antimicrobial agents and chemotherapy. 2014;58(5):2972–5.

46. Fernández-Cuenca F, Pascual A, Ribera A, Vila J, Bou G, Cisneros JM, et al. [Clonal diversity and antimicrobial susceptibility of Acinetobacter baumannii isolated in Spain. A nationwide multicenter study: GEIH-Ab project (2000)]. Enfermedades infecciosas y microbiologia clinica. 2004;22(5):267–71.

47. Fernández-Reyes M, Rodríguez-Falcón M, Chiva C, Pachón J, Andreu D, Rivas L. The cost of resistance to colistin in Acinetobacter baumannii: a proteomic perspective. Proteomics. 2009;9(6):1632–45.

48. Medeiros AA, O’Brien TF, Wacker WEC, Yulug NF. Effect of Salt Concentration on the Apparent In-Vitro Susceptibility of Pseudomonas and Other Gram-Negative Bacilli to Gentamicin. The Journal of infectious diseases. 1971;124(Supplement_1):S59–S64.

49. WHO. <WHO-PPL-Short_Summary_25Feb-ET_NM_WHO.pdfWHO Priority list.pdf>.

50. Raetz CR, Whitfield C. Lipopolysaccharide endotoxins. Annual review of biochemistry. 2002;71:635–700.

51. Bartholomew TL, Kidd TJ, Sá Pessoa J, Conde Álvarez R, Bengoechea JA. 2-Hydroxylation of <em>Acinetobacter baumannii</em> Lipid A Contributes to Virulence. Infection and immunity. 2019;87(4):e00066–19.

52. Beceiro A, Llobet E, Aranda J, Bengoechea JA, Doumith M, Hornsey M, et al. Phosphoethanolamine modification of lipid A in colistin-resistant variants of Acinetobacter baumannii mediated by the pmrAB two-component regulatory system. Antimicrobial agents and chemotherapy. 2011;55(7):3370–9.

53. Ernst RK, Moskowitz SM, Emerson JC, Kraig GM, Adams KN, Harvey MD, et al. Unique lipid a modifications in Pseudomonas aeruginosa isolated from the airways of patients with cystic fibrosis. The Journal of infectious diseases. 2007;196(7):1088–92.

54. Moskowitz SM, Ernst RK, Miller SI. PmrAB, a two-component regulatory system of Pseudomonas aeruginosa that modulates resistance to cationic antimicrobial peptides and addition of aminoarabinose to lipid A. Journal of bacteriology. 2004;186(2):575–9.

55. Llobet E, Martínez-Moliner V, Moranta D, Dahlström KM, Regueiro V, Tomás A, et al. Deciphering tissue-induced Klebsiella pneumoniae lipid A structure. Proc Natl Acad Sci U S A. 2015;112(46):E6369–78.

56. Paczosa MK, Mecsas J. <span class=“named-content genus-species” id=“named-content-1”>Klebsiella pneumoniae</span>: Going on the Offense with a Strong Defense. Microbiology and Molecular Biology Reviews. 2016;80(3):629–61.

57. Cole JN, Nizet V. Bacterial Evasion of Host Antimicrobial Peptide Defenses. Microbiol Spectr. 2016;4(1):10.1128/microbiolspec.VMBF-0006-2015.

58. Spohn R, Daruka L, Lazar V, Martins A, Vidovics F, Grezal G, et al. Integrated evolutionary analysis reveals antimicrobial peptides with limited resistance. Nature communications. 2019;10(1):4538.

59. Elliott AG, Huang JX, Neve S, Zuegg J, Edwards IA, Cain AK, et al. An amphipathic peptide with antibiotic activity against multidrug-resistant Gram-negative bacteria. Nature communications. 2020;11(1):3184.

60. Llobet E, Campos MA, Giménez P, Moranta D, Bengoechea JA. Analysis of the networks controlling the antimicrobial-peptide-dependent induction of Klebsiella pneumoniae virulence factors. Infection and immunity. 2011;79(9):3718–32.

61. Guo L, Lim KB, Poduje CM, Daniel M, Gunn JS, Hackett M, et al. Lipid A acylation and bacterial resistance against vertebrate antimicrobial peptides. Cell. 1998;95(2):189–98.

62. Caliceti P, Veronese FM. Pharmacokinetic and biodistribution properties of poly(ethylene glycol)-protein conjugates. Adv Drug Deliv Rev. 2003;55(10):1261–77.

63. Mojsoska B, Zuckermann RN, Jenssen H. Structure-Activity Relationship Study of Novel Peptoids That Mimic the Structure of Antimicrobial Peptides. Antimicrobial agents and chemotherapy. 2015;59(7):4112–20.

64. Magana M, Pushpanathan M, Santos AL, Leanse L, Fernandez M, Ioannidis A, et al. The value of antimicrobial peptides in the age of resistance. Lancet Infect Dis. 2020;20(9):e216–e30.

65. Clark DJ, Maaløe O. DNA replication and the division cycle in Escherichia coli. JMolBiol. 1967;23:99–112.

66. Campion C, Charbon G, Thomsen TT, Nielsen PE, Løbner-Olesen A. Antisense inhibition of the Escherichia coli NrdAB aerobic ribonucleotide reductase is bactericidal due to induction of DNA strand breaks. The Journal of antimicrobial chemotherapy. 2021.

67. Liang T, Leung LM, Opene B, Fondrie WE, Lee YI, Chandler CE, et al. Rapid Microbial Identification and Antibiotic Resistance Detection by Mass Spectrometric Analysis of Membrane Lipids. Analytical chemistry. 2019;91(2):1286–94.

68. Wiegand I, Hilpert K, Hancock RE. Agar and broth dilution methods to determine the minimal inhibitory concentration (MIC) of antimicrobial substances. Nature protocols. 2008;3(2):163–75.

69. Knudsen JD, Frimodt-Moller N. Animal models in bacteriology. Contrib Microbiol. 2001;9:1–14.

70. Frimodt-Moller N. The mouse peritonitis model: present and future use. J Antimicrob Chemother. 1993;31 Suppl D:55–60.

71. Agency EM. European Medicines Agency completes review of polymyxin-based medicines. 2014.

